# *Mecp2* knock-out astrocytes affect synaptogenesis by IL-6 dependent mechanisms

**DOI:** 10.1101/2023.02.08.527630

**Authors:** E. Albizzati, E. Florio, M. Breccia, C. Cabasino, D. Pozzi, E. Boda, C. Battaglia, C. De Palma, C. De Quattro, N. Landsberger, A. Frasca

**Affiliations:** Department of Medical Biotechnology and Translational Medicine, University of Milan, 20054 Segrate, Milan, Italy; Department of Biomedical Sciences, Humanitas University, 20090 Milan, Italy; IRCCS Humanitas Research Hospital, 20089 Rozzano, Milan, Italy; Department of Neuroscience Rita Levi-Montalcini, University of Turin, I-10126 Turin, Italy; Neuroscience Institute Cavalieri Ottolenghi, I-10043 Orbassano, Turin, Italy; Division of Neuroscience, IRCCS San Raffaele Scientific Institute, 20132 Milan, Italy; Department of Biotechnology, University of Verona, Cà Vignal 1, 37134 Verona, Italy

**Keywords:** Astrocytes, Synaptogenesis, Rett syndrome, Mecp2, astroglia-neuron crosstalk, Interleukin-6, paracrine molecules, non-cell autonomous mechanisms

## Abstract

Synaptic abnormalities represent a hallmark for several neurological diseases and clarification of the underlying mechanisms constitutes a crucial step towards the development of therapeutic strategies. Rett syndrome (RTT) is a rare neurodevelopmental disorder, mainly affecting females, caused by heterozygous mutations in the X-linked Methyl-CpG-Binding Protein 2 (*MECP2*) gene, leading to a deep derangement of synaptic connectivity. Although initial studies have supported the exclusive involvement of neurons, recent data have highlighted the pivotal contribution of astrocytes in RTT pathogenesis through non-cell autonomous mechanisms. Since astrocytes regulate synaptogenesis by releasing multiple molecules, we investigated the influence of soluble factors secreted by *Mecp2* KO astrocytes on synaptic density. We found that *Mecp2* deficiency in astrocytes negatively affects their ability to support synapse formation by releasing synaptotoxic molecules, among which we identified interleukin-6 (IL-6). Notably, aberrant IL-6 expression exclusively emerges from a dysfunctional astrocyte-neuron crosstalk, and blocking IL-6 activity prevents synaptic alterations.

## Introduction

Astrocytes represent one of the most abundant classes of glial cells in the mammalian brain and play a crucial role for proper health and function of the central nervous system (CNS), providing metabolic and trophic support to neurons^1^. Accordingly, astrocyte-neuron interactions are fundamental for synaptic development across different brain regions since early embryonic stages to adulthood^2,3^. By secreting bioactive proteins, such as neurotrophins, synaptogenic cues, cytokines and chemokines, astrocytes finely promote synaptic formation, functional maturation and refinement^3–6^. Consequently, altered secretion of these molecules can contribute to synaptic defects, as evidenced in many neurodevelopmental, neuropsychiatric and neurodegenerative disorders^7–10^. Rett syndrome (RTT) is a rare neurodevelopmental disease caused in the vast majority of cases by mutations in the X-linked Methyl-CpG-binding Protein 2 (*MECP2*)^11^. Besides neurons, which suffer from severe morphological, functional and synaptic defects^12–15^, recent data indicated a role for astrocytes in the disease, reporting that *Mecp2* mutant astrocytes and their conditioned medium exert a negative effect on neuronal development, decreasing dendritic outgrowth and affecting overall maturation^16–18^. Accordingly, the solely loss of *Mecp2* in astrocytes is sufficient to cause pathological alterations, while rescuing *Mecp2* expression specifically in these cells greatly ameliorates RTT symptoms and leads to synaptic improvements, suggesting the possibility to target astrocytes for therapeutic purpose^17^. Two proteomic studies analyzed deregulated proteins secreted by *Mecp2* KO astrocytes to disclose molecules influencing dendritic maturation. Ehinger and colleagues examined the secretome of KO cortical astrocytes isolated from P1-2 pups, revealing the decrease of Lcn2 and Lgals3, that, when added to null neurons, effectively improved dendritic arborization^19^. More recently, Caldwell and collaborators analysed the proteome of culture medium from KO cortical astrocytes isolated at P7 by immunopanning, reporting the alteration of several proteins, including an increase of Igfbp2 and BMP6, that were highlighted as molecular candidates involved in the occurrence of neuronal defects^20^. However, the above-mentioned studies analysed the secretome of astrocytes cultured alone, without considering their crosstalk with neurons, which strongly affects molecular and functional properties of both cell populations^21–23^, an issue particularly relevant for RTT, that is characterized by mosaic interactions between cells expressing either the wildtype or mutant *MECP2* allele. Thus, in our study, we explored whether and how *Mecp2* KO astrocytes might alter synaptogenesis in neurons, investigating the astrocyte-neuron crosstalk in a co-culture system. We report that KO astrocytes secrete molecules that affect synaptogenesis in WT neurons and transcriptomic analyses suggested the involvement of genes related to inflammatory response. Molecular investigations revealed an increase of Interleukin-6 (IL-6) levels in co-culture medium and, accordingly, of its transcription in KO astrocytes. We thus confirmed the causative role of this cytokine for the induced synaptic impairments. Interestingly, excessive secretion of IL-6 exclusively occurs when astrocyte-neuronal crosstalk is preserved and lacks when KO astrocytes are cultured alone.

All in all, our study identifies a novel pathogenic mechanism triggered by the communication between neurons and *Mecp2* KO astrocytes, that leads to synaptic alterations and might provide a novel therapeutic target for RTT and other *MECP2*-related disorders.

## Materials and methods

### Animals

The *Mecp2^tm1.1^* Bird mouse strain was originally purchased from the Jackson Laboratories and then backcrossed and maintained on a clean CD1 background^44^. These mice recapitulate the typical phenotype of C57BL/6 mice, with the advantage of having a larger progeny and minor risk of litter cannibalization. Mouse genotype was determined by PCR on genomic DNA purified from ear punch biopsies, and both KO male and HET female animals were used for different experiments. Mice were housed in a temperature- and humidity-controlled environment in a 12-h light/12-h dark cycle with food and water *ad libitum*. All procedures were performed in accordance with the European Union Communities Council Directive (2010/63/EU) and Italian laws (D.L.26/2014). Protocols were approved by the Italian Council on Animal Care in accordance with the Italian law (Italian Government decree No. 210/2017 and 187/2022).

### Primary cultures of neurons

Primary cortical neurons were prepared from WT mouse embryos generated by mating WT females with WT male mice. The day of vaginal plug was considered E0.5 and primary neurons were prepared from E15.5 embryos to avoid glial contamination. Embryos were sacrificed by decapitation, brains were removed under a microscope and rapidly immersed in ice-cold HBSS. Meninges were gently removed, and cerebral cortex from both hemispheres was rapidly dissected into small pieces and maintained in cold HBSS until tissue dissociation. Tissues were washed in HBSS, incubated with 0.25% trypsin/EDTA for 7 min at 37°C and the digestion was blocked with dissociation medium (DMEM HG containing 10% FBS and 1% Pen/Strep). Then, cortices were accurately washed and mechanically dissociated by gently pipetting. Cells were counted (Countess Automated Cell Counter, ThermoFisher) and, depending on experimental needs, neurons were seeded on poly-D-lysine-coated plates (0.1 mg/mL), poly-D-lysine-coated glass coverslips (1 mg/mL) or directly on astrocytes, at the density described below.

### Primary cultures of astrocytes

Primary cultures of astrocytes were prepared from cerebral tissue of P1-P3 WT and *Mecp2* null mice (P3 only in case of cerebellar astrocytes), generated by mating *Mecp2* HET females with WT male mice. Pups were decapitated and brains immediately collected on ice-cold HBSS. Meninges were carefully removed and, depending on the experiment, cortices, hippocampi or cerebella were isolated from both hemispheres and immersed in HBSS containing 10 mM HEPES, 4 mM Na_2_HCO_3_ and 1% P/S until dissociation. Mouse genotype was determined by PCR on genomic DNA purified with Phire animal tissue direct PCR kit. Tissues were then incubated in 0.25% trypsin/EDTA for 30 min at 37°C, mechanically dissociated in astrocyte culture medium (DMEM HG containing pyruvate plus 1:1 Ham’s F-10, 10% FBS and 1% Pen/Strep) with a glass Pasteur and filtered through cell strainers of 40 μm pore size to obtain a single cell suspension. The resulting cells were centrifuged at 1,500 g for 7 min, re-suspended in culture medium, and plated in poly-D-lysine-coated 75 cm^2^ flasks (15 μg/mL). At DIV4, flasks were shaked in astrocyte culture medium containing 10 mM HEPES at 200 rpm for 8 h at 37°C to eliminate other cell types, and the medium was replaced with fresh culture medium. Cells were incubated in a humidified incubator at 37°C and 5% CO_2_ and culture medium was refreshed every 3-4 days. When astrocytes reached confluence (DIV12-DIV15), they were washed in D-PBS to remove death cells or cell debris, and then detached by 0.25% trypsin/EDTA diluted in HBSS (1:2 ratio) and counted in a Bürker chamber. Depending on experimental needs, they were seeded on poly-D-lysine-coated plates (15 μg/mL), poly-D-lysine-coated glass coverslips (1 mg/mL) or transwell membrane inserts, at the density described below.

### Isolation of mouse cortical astrocytes

To perform gene expression analysis on astrocytes acutely sorted from P7 *Mecp2* HET mice and the corresponding WT female littermates, animals were sacrificed by rapid decapitation and brains collected in ice-cold HBSS. Meninges were carefully removed and cortices from both hemispheres were isolated and immersed in HBSS containing 10 mM HEPES, 4 mM NaHCO3 and 1% P/S until dissociation. Tissues were then incubated in 0.05% trypsin/EDTA containing DNase I (1:1,000) for 5 min at 37°C, mechanically dissociated by gently pipetting and filtered through cell strainers of 40 μm pore size adding two volumes of astrocyte culture medium (DMEM plus 1:1 Ham’s F-10, 10% FBS and 1% P/S). Cell suspensions were then centrifuged at 1,000 rpm for 10 minutes at 4°C. Positive selection of astrocytes was performed by magnetic labelling of ACSA1-positive cells, by using a biotin-conjugate anti-ACSA1 antibody and the following labelling with streptavidin-coated magnetic beads. Labelled cells were retained by ferromagnetic columns placed in a magnetic field^45^. Finally, cells were directly lysed in 1 mL PureZOL and stored at −80°C until RNA extraction.

### Astrocyte-neuron co-cultures

#### In contact co-cultures

Astrocytes (30,000 cells) were seeded on glass coverslips and after 2-4 days, neurons were added on the top of astrocyte monolayers, at low density (10,000 cells/well). Cells were cultured, during the first 48 hours, in a 5% FBS co-culture medium (Neurobasal medium containing 5% FBS, 2% B27, 1% L-Glutamine, 1% P/S), then in a medium containing lower serum concentration (Neurobasal medium containing 2,5% FBS, 2% B27, 1% L-Glutamine, 1% P/S) for the entire duration of the experiment, corresponding to 14 days for neurons. Co-cultures were filled with fresh medium at DIV7 for a third of the original volume.

#### Transwell co-cultures

Astrocytes were seeded at a density of 8,000 cells/transwell membrane inserts for 24-well plates or of 150,000 cells/transwell membrane for 6-well plates. Neurons (derived from 3 different embryos per experiment) were seeded at a density of 30,000 cells/glass coverslip for 24-well plates or 200,000 cells/well for 6-well plates. Inserts with astrocytes were maintained in their culture medium for 4 days, with a change of the medium after 2 days to remove cells that did not attach. Then, medium was replaced with neuron culture medium and inserts were carefully transferred above neurons ~1 hour after seeding. The transwell-based co-culture was maintained until DIV14. At DIV7, both neurons and astrocytes were filled with fresh neuron culture medium for a third of the original volume, except when treated with IL-6 neutralizing antibody (see the specific section).

### Astrocyte Conditioned Medium (ACM) preparation and treatment

For ACM preparation, astrocytes were seeded at a density of 100,000 cells/well in 6-well plates. When confluent, they underwent starvation by removing FBS from medium. Briefly, after washing cells twice in DMEM (a rapid wash, followed by an incubation of 90 minutes), the medium was replaced by neuronal culture medium without B27 (Neurobasal containing 1% L-Glutamine and 1% P/S). After 48 hours, ACM was collected and centrifuged at 1,200 rpm for 5 minutes at 4°C to remove cell debris. A cocktail of proteases inhibitor was added 1:1,000 to each sample and ACM was stored at −80°C until use. For ACM treatment, neurons (30,000 cells/coverslip) were incubated with pre-warmed (37°C) ACM at DIV13 for 24 hours. Heat inactivation of ACM was performed by boiling it at 95°C for 5 minutes and slowly cooling it down to 37°C prior to neuronal treatment. Neuronal culture medium was added as control on neurons (NT).

### Treatment of co-cultures with neutralizing IL-6 antibody

Transwell based co-cultures were treated at DIV5 and DIV12 with either neutralizing antibody for IL-6 (0.5 mg/ml) or an isotypic antibody Rat IgG1 (1 mg/ml), diluted 1:500 and 1:1,000, respectively, in the final volume of culture medium (Neurobasal containing 2% B27, 1% L-Glutamine, 1% P/S). Untreated co-cultures were filled with an equal volume of culture medium. In addition, at DIV5, transwell inserts were filled with either 50 μL (when in 24-well plates) or 100 μL (when in 6-well plates) of medium.

### RNA Sequencing and bioinformatic analyses

Primary neurons were lysed in PureZOL and RNA was extracted following manufactures instructions. Samples were incubated with DNase at 37°C for 15 minutes to remove eventual genomic DNA contaminations. Then, enzymatic reaction was inactivated in PureZOL and a second RNA extraction was performed.

Generation of RNA-Seq data was performed by GENARTIS srl (Verona, Italy), as follows. RNA purity was measured at a NanoDrop Spectrophotometer (ThermoFisher Scientific) and RNA integrity was assessed using the RNA 6000 Nano Kit on a Bioanalyzer (Agilent Technologies). All samples showed an RNA integrity number (RIN)>9. RNA samples were quantified using the Qubit RNA BR Assay Kit (ThermoFisher Scientific). RNA-Seq libraries were generated using the TruSeq stranded mRNA kit (Illumina) from 400 ng of RNA samples, after poly(A) capture and according to manufacturer’s instructions. Quality and size of RNA-Seq libraries were assessed by capillary electrophoretic analysis with the Agilent 4200 Tape station (Agilent Technologies). Libraries were quantified by real-time PCR against a standard curve with the KAPA Library Quantification Kit (KapaBiosystems, Wilmington, MA, USA). Libraries were pooled at equimolar concentration and sequenced on a NovaSeq6000 (Illumina) generating >20 million fragments in 150PE mode for each sample. Sequencing read trimming and quality of reads were assessed using FastQC software (http://www.bioinformatics.babraham.ac.uk/projects/fastqc/). Starting from raw FASTQ files, the first 10 nt of read2 were trimmed as they presented a lower quality than expected (Q<30). In addition, reads with more than 10% of undetermined bases or more than 50 bases with a quality score <7 were discarded. Reads were then clipped from adapter sequences using Scythe software (v0.991) (https://github.com/vsbuffalo/scythe), and low-quality ends (Q score <20 on a 10-nt window) were trimmed with Sickle (v1.33) (https://github.com/vsbuffalo/sickle). Filtered reads were aligned to the Mouse reference genome GRCm38 (Ensembl release 102) using STAR (v2.7.6a) with default parameters and quantMode TranscriptomeSAM option that output alignments translated into transcript coordinates. After reads mapping, the distribution of reads across known gene features, such as exons (CDS, 5’UTR, 3’UTR), introns and intergenic regions, was verified using the script read_distribution.py provided by RSeQC package (v3.0.1). Read counts on genes were quantified using RSEM (v.1.3.3) and Mouse Ensembl release 102 annotation. Genes-level abundance, estimated counts and gene length obtained with RSEM were summarized into a matrix using the R package tximport (v1.18.0) and, subsequently, the differential expression analysis was performed with DESeq2 (v1.30.0) integrating the “day of sample preparation” as variable in the model. To generate more accurate Log2 FoldChange estimates, the shrinkage of the Log2 FoldChange was performed applying the apeglm method.

Gene Ontology (GO) enrichment analysis was performed using clusterProfiler, an R Package for comparing biological themes among gene clusters^46^ (Bioconductor version: Release (3.18.0)). The function simplify was used to remove redundancy of enriched GO terms. Differentially expressed genes (DEGs) with p.adj<0.1 were included in the analysis, that was performed on all the 3 comparison groups. FDR adjusted p-value (q-value) <0.05 was used as a threshold and GO terms fulfilling this condition were defined as significantly enriched.

Preranked Gene Set Enrichment Analysis (GSEA)^47^ (version 4.1.0, the Broad Institute of MIT and Harvard; https://www.gsea-msigdb.org/gsea/downloads.jsp) was performed on a pre-ordered gene list ranked according to the log2 fold changes from DESeq2, between WT neurons cultured with *Mecp2* KO astrocytes and WT neurons cultured with WT astrocytes. GSEA calculated a Normalized Enrichment Score (NES) of gene sets present in the MsigDB 7.2 (Molecular Signatures Database; https://www.gsea-msigdb.org/gsea/msigdb), with 1,000 permutations set to generate a null distribution for enrichment score. ‘c5.go.bp.v7.2.symbols.gmt’ was the gene set database used for enrichment analysis and FDR q-value<0.05 was defined as the cut-off criteria for significance. Metascape analyses were performed using the Metascape platform^25^ (version 3.5; https://metascape.org) and giving in input gene lists of upregulated DEGs with p.adj<0.1 of comparisons ‘+aWT’ *versus* CTRL (3239 DEGs, **Table S4**) and ‘+aKO’ *versus* CTRL (3808 DEGs, **Table S5**).

### Quantitative Reverse Transcriptase PCR (qRT-PCR)

Total RNA was extracted from astrocytes cultured on transwell membrane inserts or on 6-well plates, and from astrocytes sorted from P7 mouse cortices using PureZOL. RNA was quantified using a NanoDrop 1000 spectrophotometer (ThermoFisher Scientific) and its integrity verified by agarose gel electrophoresis. cDNA was synthesized using the RT^2^ First Strand Kit according to the manufacturer’s instructions and used as template for quantitative RT-qPCR with SYBR Green Master Mix and a QuantStudio 5 Real-Time PCR System (ThermoFisher Scientific). Due to the low amount of RNA extracted from astrocytes on transwell, a preamplification step was performed using SsoAdvanced™ PreAmp Supermix prior to qPCR, following manufacturer’s instructions. Melting curve showed a single product peak, indicating good product specificity. The best housekeeping gene was selected for each comparison (*Rpl13* for astrocytes cultured alone and sorted from cortex and *Ywhaz* for astrocytes on transwell), and fold change in gene expression was calculated using the 2^(-Delta Ct) method.

### Immunofluorescence

Neurons seeded on glass coverslips were fixed in 4% paraformaldehyde (PFA) plus 10% sucrose for 8 minutes at RT and, after three washes in 10mM PBS, they were stored in 10 mM PBS containing 0.1% sodium azide at 4°C until staining. To assess the efficiency of MACS sorting, 10 μl of cell suspension was transferred on glass coverslips and cells stained for astrocytic marker. First, cells were washed in 10 mM PBS and then permeabilized in PBS containing 0.2% Triton X-100 for 3 minutes on ice. After three washes in PBS containing 0.2% BSA, blocking solution (4% BSA in PBS) was added for 15 minutes before incubating the primary antibodies (Synapsin1/2 1:500, Shank2 1:300, MAP2 1:1000 on neurons; GFAP 1:500 for astrocytes), diluted in PBS containing 0.2% BSA overnight (4°C). Then, cells were washed in PBS containing 0.2% BSA before incubation (1 hour at RT) with Alexa Fluor secondary antibodies, diluted 1:500 in PBS containing 0.2% BSA. Several washes in 0.2% BSA in PBS were then performed and nuclei were stained with 1:1,000 DAPI in PBS solution for 10 minutes at RT. Finally, neurons were washed in 10mM PBS before mounting the glass coverslips on glass microscope slides with Fluoromount Aqueous Mounting Medium.

### Microscopy and image analysis

#### Analysis of synaptic markers and colocalization

Z-stack images of synaptic puncta (1,024 × 1,024 pixel resolution) were acquired either at 1× digital zoom using a SR Apo TIRF 100X oil-immersion objective mounted on a Nikon Ti2 microscope equipped with an A1+ laser-scanning confocal system (16-bit grayscale depth images) or at 1.4× digital zoom using a 63× oil-immersion objective mounted on a Leica DMI3000B microscope equipped with a Sp5 laser-scanning confocal system (8-bit grayscale depth images), depending on the experiment. Acquisition of synaptic puncta was performed using a step size of 0.3 μm. For each dataset acquisition, parameters (offset background, digital gain, and laser intensity) were maintained constant among different experimental groups. By Fiji software, maximum intensity projection images were converted to binary images and processed with a fixed threshold for each channel acquired. Puncta density was calculated by counting only puncta lying along manually selected ROIs within 20 μm of 3 primary branches/neuron. Only puncta with a minimum size of 0.16 μm^2^ were counted using *Analyze Particles^48^*. To assess puncta co-localization of pre- and post-synaptic markers, the Fiji Plugin *Colocalization highlighter* was run on each Z-stack image acquired. Co-localized puncta were quantified in manually selected ROIs of the binary mask created from the maximum intensity projection. Only puncta with a minimum size of 0.1 μm^2^ were counted. Sample size of neurons analysed per experimental group was assessed based on previous work^49^.

#### Western blot

To perform Western blots on neurons, cells were rapidly washed with DPBS at RT and then lysed by a cell scraper in sample buffer containing 2-Mercaptoethanol (1:10 v/v), protease (1:200 v/v) and phosphatase (1:10 v/v) inhibitors and sonicated at 30 Hz for 5 seconds. After denaturation at 70°C for 5 minutes, equal volume of each sample (15 μl) was resolved on 4–15% Criterion TGX Precast Protein Gels. Before transfer, a Stain-Free gel image was acquired using a UV-transilluminator (ESSENTIAL V6 System, UVITEC Ltd, UK) and used for relative quantification. Proteins were blotted on a nitrocellulose membrane using the Trans-blot SD (Bio-Rad) semidry apparatus and ponceau S staining was used to verify proper protein transfer. Membranes were then incubated at RT for 1 hr in blocking solution (Tris-buffered saline containing 0.5% Tween-20 (TBST) and 5% nonfat milk or 5% BSA) before adding primary antibodies at the proper dilution: anti-NF-kB p65 (1:500 in 5% milk-TBST); anti-phospho-Stat3 (1:1,000 in 5% BSA-TBST); anti-Stat3 (1:2,500 in 5% BSA-TBST); anti-Gapdh (1:10,000 in 5% milk-TBST). After 3 washes in TBST, blots were incubated with the appropriate HRP-conjugated secondary antibody diluted 1:10,000 in 5% milk or 5% BSA-TBST. Immunocomplexes were visualized using the ECL substrates kits from Cyanagen and imaged on ALLIANCE MINI HD9 system (UVITEC Ltd, UK). Quantification of bands was performed using the Uvitec Nine Alliance Software, with the relative density of bands corrected on background signal. Phospho-Stat3 levels were normalized to total Stat3 levels, while p65 signal was normalized on Gapdh. Percentages plotted on graphs were calculated by normalizing all conditions to the control group.

#### Cytokines quantification

Cytokines were detected in culture media by FRACTAL Unit (Flow cytometry Resource, Advanced Cytometry Technical Applications Laboratory, San Raffaele Scientific Institute, Milan) with two LEGENDplex assay kits: LEGENDplex Mouse Cytokine Release Syndrome Panel and LEGENDplex Mouse Cytokine Panel 2 according to the manufacturer’s instructions. Cell culture media were used undiluted. Samples were acquired using a BD FACSCantoTM II Cell Analyzer (BD Biosciences) using BD FACS Diva software and equipped with three lasers: blue (488 nm), red (633 nm) and violet (405). FCS files were analyzed by using LEGENDplex^™^ Data Analysis Software (BioLegend) according to the manufacturer’s instructions, obtaining the absolute concentration for each molecule.

#### Quantification and statistical analysis

Data are expressed as mean±SEM, except for violin plots in which the bar indicates the median with interquartile range. Replicates are indicated in figure legends. Before any statistical analysis, normality distribution was evaluated for each dataset by D’Agostino and Pearson test and outliers were assessed by ROUT test (Q = 1%) or Grubb’s test (a = 0.05%). Unpaired Student’s t-tests or Mann–Whitney tests were used for the comparisons of two groups, in accordance with data distribution. To assess the effect of ACM treatment on WT neurons, Kruskal–Wallis test followed by Dunn’s post hoc test was applied. Statistical significance for multiple group comparisons of co-culture treatments was determined by two-way ANOVA, followed by Tukey’s post hoc test. All statistical analyses were performed using Prism 8 (GraphPad Software, La Jolla, CA, United States). A p-value <0.05 was considered significant. *p<0.05, **p<0.01, ***p<0.001.

## Results

### Molecules secreted by *Mecp2* KO astrocytes affect synaptogenesis in WT neurons

Since astrocytes play key roles in synapse formation and RTT pathogenesis^3,16,17^, we exploited a transwell-based co-culture between WT neurons and *Mecp2* KO astrocytes to investigate the consequence of *Mecp2* deficiency in astrocytes on synaptogenesis. WT-WT co-cultures were used as control. The impact on synaptogenesis was assessed by analysing the density of pre- and post-synaptic puncta, and their colocalization, as an index of functional synapses (**Figure 1A**). WT neurons matured under the influence of KO cortical astrocytes displayed reduced number of both pre- and post-synaptic puncta as well as of colocalized puncta, compared to control (**Figure 1B, C**), with no change in size (**Figure S1**). Of note, KO astrocytes cultured in close contact with WT neurons caused reduction of pre-synaptic puncta density, as well as of puncta area (**Figure S2**), thus suggesting that other factors beyond secreted molecules might alter the synaptic phenotype.

**Figure 1.**
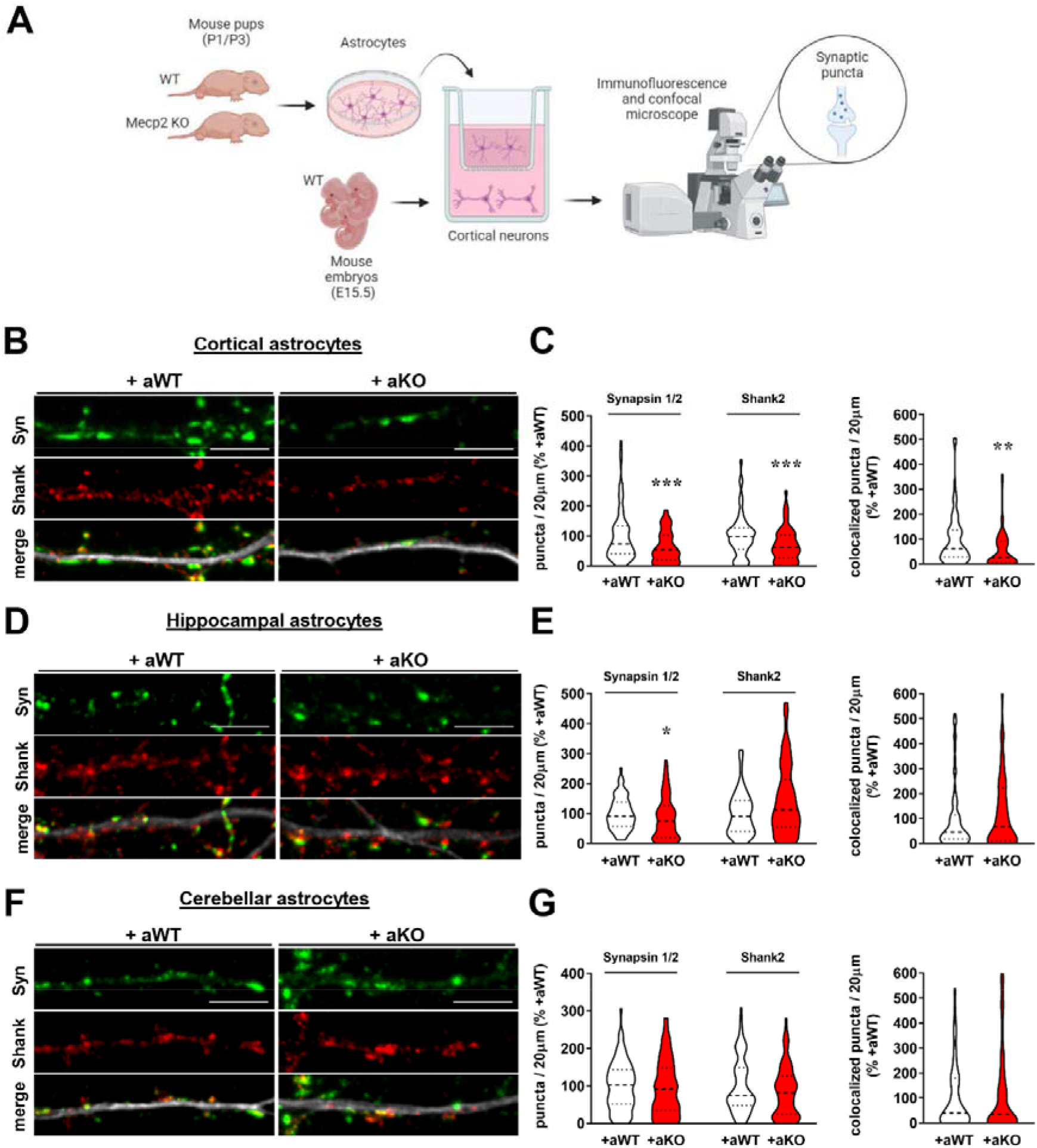
Soluble factors secreted by *Mecp2* KO astrocytes affect synaptogenesis, with cortical astrocytes showing the most detrimental synaptogenic effects. **(A)** Experimental design overview, created with BioRender.com. **(B,D,F)** Representative images of primary branches from WT neurons (DIV14) immunostained for Synapsin1/2 (green), Shank2 (red) and their merge with MAP2 (white), in co-culture with cortical (B), hippocampal (D) and cerebellar astrocytes (F). Scale bar = 5 μm. **(C,E,G)** Violin plots indicate the median (dashed line) and 25^th^ and 75^th^ percentiles (dotted lines) of Synapsin1/2, Shank2 and colocalized puncta density. Values for puncta number are expressed as percentages compared to WT-WT co-cultures (set at 100%). *p<0.05, **p<0.01, ***p<0.001 by Mann Whitney test. Analyses were performed on n>60 neurons (in C), on n>66 neurons (in E) and on n>46 neurons (in G) per experimental group from N>7 (in C and E) or N>5 (in G) biological replicates. All data derived from at least 2 independent experiments.

According to the heterogeneous properties of astrocytes depending on cerebral region, we assessed whether *Mecp2* deficiency diversely affects their synaptogenic potential when derived from brain regions other than cortex^24^. Interestingly, WT neurons cultured with hippocampal KO astrocytes exhibited a selective impairment of the density of pre-synaptic puncta, with no effect on post-synapses and colocalization (**Figure 1D, E**), whilst cerebellar astrocytes did not cause any synaptic defect (**Figure 1F, G**).

These results demonstrate that molecules secreted by *Mecp2* KO cortical astrocytes impair synaptogenesis, therefore supporting a non-cell autonomous influence on WT neurons. The release of neurotoxic factors and/or defective secretion of synaptogenic molecules might be involved. Comparing synaptogenic potentials of astrocytes from cortex, hippocampus and cerebellum, we report a heterogeneous response to *Mecp2* loss, indicating cortical astrocytes as the most affected cells.

### Transcriptional profile of WT neurons confirms the detrimental action of *Mecp2* KO astrocytes on synapse formation, highlighting the role of pro-inflammatory cues

To gain insights into the effects mediated by astrocytes on neurons, we performed bulk RNA-Seq on neurons cultured with astrocytes, when seeded on transwell inserts (**Figure 2**). Transcriptomic analysis was conducted on three experimental groups: WT neurons cultured alone (“CTRL”), WT neurons cultured with WT astrocytes (“+aWT”) and WT neurons cultured with *Mecp2* KO astrocytes (“+aKO”) (**Figure 2A**). Transcriptional profiles of +aWT or +aKO groups were compared to each other (+aKO *versus* +aWT) and to that of CTRL (+aWT *versus* CTRL; +aKO *versus* CTRL). Principal Component Analysis (PCA) of whole transcriptome expression highlighted that both co-culture systems clustered together with respect to CTRL (**Figure S3A**), confirming that neurons dramatically change their transcriptional profiles when cultivated with astrocytes. Indeed, both co-cultures showed a great number of differentially expressed genes (DEGs) when compared to CTRL, whereas few significant DEGs emerged from the comparison of the two co-cultures (**Figure 2B** and **Figure S3B**).

**Figure 2.**
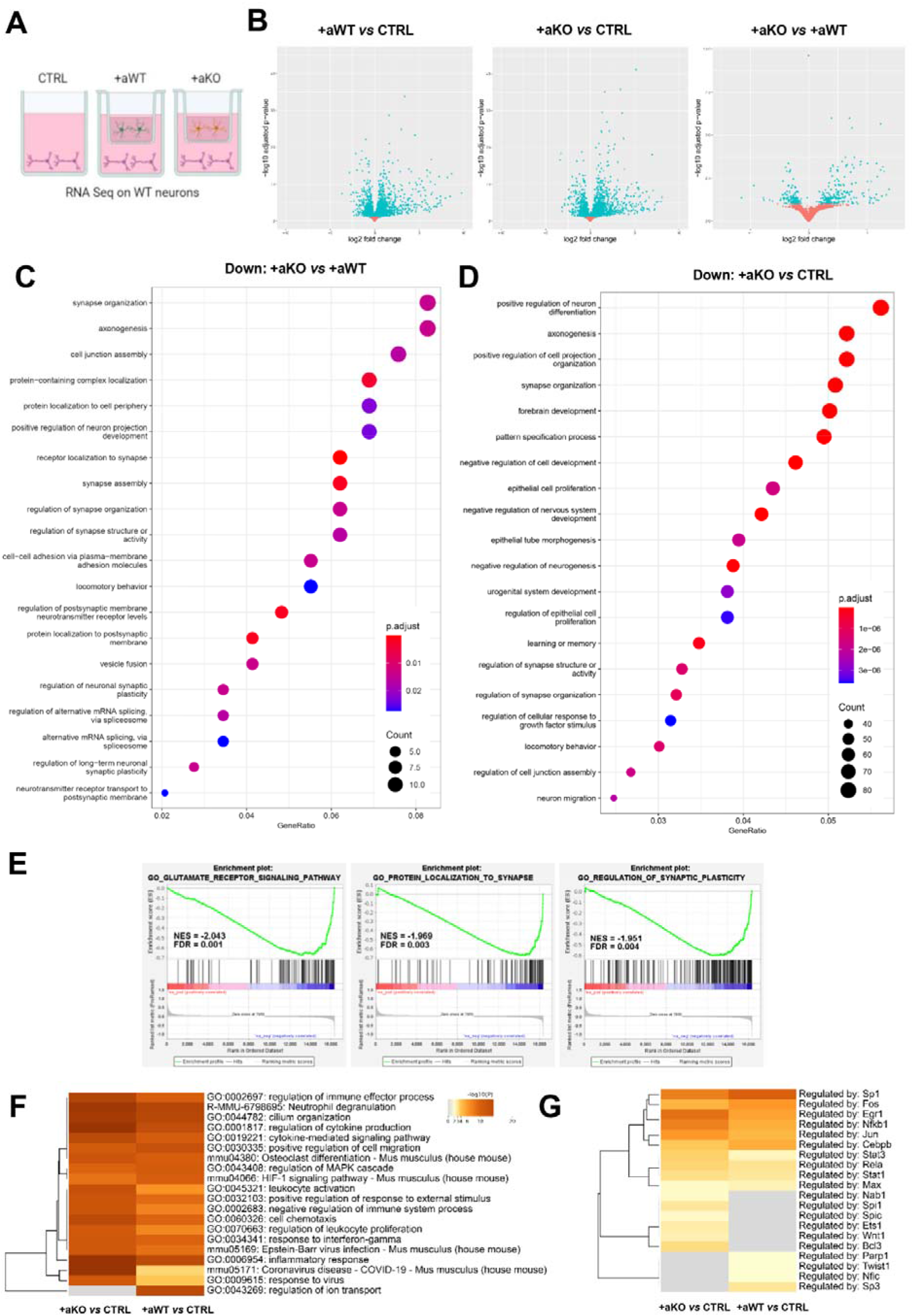
*Mecp2* KO astrocytes affect the expression of genes involved in synaptic maturation and inflammatory responses in WT co-cultured neurons. **(A)** Experimental groups, created by BioRender.com, included in RNA-seq analysis: neurons cultured alone (CTRL) (n=5) and neurons matured in co-culture with WT astrocytes (+aWT) (n=7) or *Mecp2* KO astrocytes (+aKO) (n=8). Replicate samples were derived from 2 independent experiments. **(B)** Volcano plots of DEGs with p.adj<0.1 from the different comparisons, plotted as log2FC against −log_10_ adjusted p-value. Each dot represents a gene: top-right sector highlights genes that are significantly upregulated, top-left sector genes that are significantly downregulated, with p.adj<0.1. **(C,D)** Enrichment analysis of downregulated DEGs at p.adj<0.1 in +aKO *vs* +aWT (C) and +aKO *vs* CTRL (D) comparisons (see Table S1 and S2, respectively), showing the top 20 significant GO terms (biological processes). Size and colour of dots represent number of genes associated with each term and FDR adjusted p-value, respectively, according to the scale indicated in the figure. **(E)** Enrichment plots from preranked GSEA of the comparison +aKO *vs* +aWT. Three of the top ten negatively correlated gene sets are reported, together with their Normalized Enrichment Score (NES) and significance (FDR) (see Table S3 for complete results). **(F,G)** Metascape analyses of upregulated DEGs (see Table S4 and S5) from +aWT *vs* CTRL and +aKO *vs* CTRL. Top 20 most represented biological processes together with their statistical value (represented as − log10 p-value) (F) and enrichment analysis of transcriptional regulators using the TRRUST database (G).

In order to identify the main biological processes engaged in the different conditions, Gene Ontology (GO) enrichment analysis was performed separately on downregulated or upregulated DEGs. Interestingly, comparing the downregulated DEGs of +aKO *versus* +aWT, processes related to synapse organization, activity and axonogenesis emerged as the most affected ones (**Figure 2C** and **Table S1**). These results greatly confirmed the immunofluorescence data, highlighting the impact of *Mecp2* deficient astrocytes on proper synaptogenesis and neuronal maturation. Of note, downregulation of similar pathways also occurred by comparing neurons cultured with KO astrocytes to CTRL, thus indicating the presence of synaptotoxic paracrine signals rather than the lack of trophic factors (**Figure 2D and Table S2**).

To better investigate the overall molecular pathways affected in WT neurons by KO astrocytes, a preranked Gene Set Enrichment Analysis (GSEA) was conducted on the comparison +aKO *versus* +aWT. The analysis yielded several negatively correlated gene sets and only few positively correlated gene sets (NES<0 or >0, respectively, combined with FDR q<0.05, **Table S3**). Among the top ten negatively correlated gene sets, we found *‘glutamate receptor signaling pathway’*, *‘protein localization to synapse’* and *‘regulation of synaptic plasticity’,* corroborating synaptic alterations emerged from immunostaining experiments (**Figure 2E**).

Since the direct comparison of the two co-cultures did not provide any suggestions for the onset of synaptic alterations, we focused on the comparison of +aWT *vs* CTRL and +aKO *vs* CTRL, by exploring the upregulated pathways derived from the GO analyses (**Figure S3C,D**). Interestingly, the analysis of +aWT *vs* CTRL unmasked pathways related to ion transport, glutamatergic transmission and synaptic organization, that were not present in +aKO *vs* CTRL, reinforcing the inability of KO astrocytes to support synaptic maturation (**Table S4,5**). Further, many biological processes associated with cytokines production/signalling and immune response were enriched in both GO analyses. To better extrapolate the entity of inflammatory pathways, we exploited Metascape resources on upregulated DEGs^25^, finding that responses to cytokines and inflammatory stimuli were more represented in the +aKO *vs* CTRL with respect to the +aWT *vs* CTRL comparison (**Figure 2F,S4 and Table S4,5**). Moreover, enrichment analysis of transcriptional regulators using the TRRUST database confirmed a stronger activation of transcriptional factors involved in inflammatory response (such as NFkB1, Jun, Stat3, Stat1, Spi1, Ets1, Wnt1, Bcl3) in the +aKO *vs* CTRL comparison (**Figure 2G**).

Overall, transcriptional analyses supported the occurrence of synaptic defects induced by *Mecp2* KO astrocytes and indicated an increased inflammatory response as candidate molecular mechanism.

### Aberrant IL-6 secretion by *Mecp2* KO astrocytes causes synaptic alterations

To confirm the ability of KO astrocytes to activate inflammatory pathways in neurons, the protein expression of the inflammatory trigger p65 subunit of NF-kB was evaluated in neurons exposed to either WT or KO astrocytes. Western blot analysis indicated that the p65 subunit was significantly increased in neurons cultured with KO astrocytes (**Figure 3A**). In parallel, we measured mRNA levels of a panel of pro-inflammatory molecules in KO astrocytes cultured on transwell inserts and their protein products’ concentration in the co-culture medium. qRT-PCR showed a strong upregulation of *IL-1β*, *IL-6* and *CXCL12* in KO astrocytes. Conversely, the expression of *HMGB1,* a trigger of inflammation in many neurodegenerative diseases, was slightly reduced (**Figure 3B**). Protein levels of the secreted cytokines and chemokines were measured by a cytometric bead-based immunoassay platform, finding a 3-fold increase in IL-6 concentration in the medium derived from co-cultures with KO astrocytes, and a slight, but not statistically significant, increase of TNFα, CCL2, CCL3 and CCL4 (**Figure 3C**).

**Figure 3.**
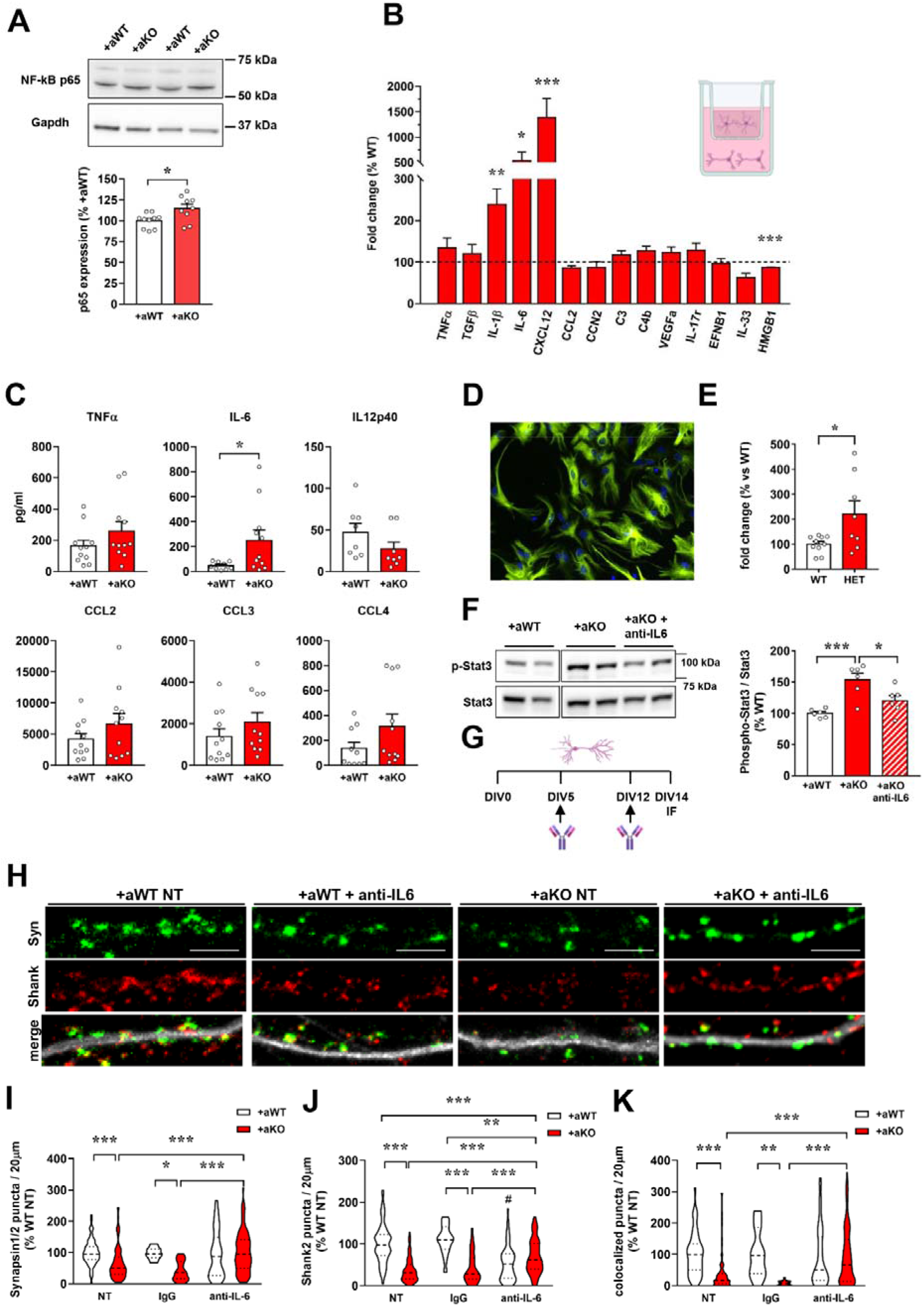
*Mecp2* KO astrocytes aberrantly secrete IL-6, which causes synaptic defects in WT neurons. **(A)** Western blot analysis of p65 protein levels in WT neurons (DIV14) cultured with *Mecp2* KO or WT astrocytes (n=10 +aWT and n=9 +aKO). Data are represented as mean±SEM and expressed as percentage of +aWT condition; representative bands for p65 and Gapdh are reported. Samples derived from 3 independent experiments. **(B)** The graph depicts the mRNA expression of selected astrocyte genes in *Mecp2* KO cortical astrocytes cultured for 14 days with WT neurons. Data are expressed as percentages of +aWT condition (n=10) and represented as mean±SEM. *p<0.05, **p<0.01, ***p<0.001 by Student’s t-test or Mann–Whitney test in accordance with data distribution. Samples derived from at least 3 independent experiments. **(C)** Cytokines’ quantification in the co-culture medium. Data are represented as mean±SEM and expressed as percentage of +aWT condition (n=11). *p<0.05 by Student’s t-test. Samples derived from at least 3 independent experiments. **(D)** Immunofluorescence staining for GFAP in acutely sorted astrocytes from P7 cortices. **(E)** The graph shows the mRNA expression of IL-6 in HET cortical astrocytes, compared to WT astrocytes. Data, expressed as percentage of WT samples, are reported as mean±SEM. Samples (n=10 WT and n=9 HET) derived from 3 independent litters. *p<0.05 by Student’s t-test. **(F)** Western blot analysis of phospho-Stat3 over Stat3 protein levels in WT neurons at DIV14 in culture with astrocytes and treated or not with anti-IL-6 antibody, as indicated in G (created with BioRender.com). Data are represented as mean±SEM and expressed as percentage of +aWT (n=12); representative western blot bands are reported. *p<0.05; ***p<0.001 by 1-way ANOVA followed by Tukey’s post hoc test. Samples derived from at least 3 independent experiments. **(H)** Representative images of primary branches from WT neurons (DIV14) immunostained for Synapsin1/2 (green), Shank2 (red) and their merge with MAP2 (white). Scale bar = 5 μm. **(I-K)** Violin plots indicate the median (dashed line) and 25^th^ and 75^th^ percentiles (dotted lines) of Synapsin1/2 (I), Shank2 (J) and colocalized puncta number (K) in neurons co-cultured with WT (+aWT) or KO (+aKO) astrocytes seeded on transwell inserts. Values for puncta number are expressed as percentages of +aWT untreated condition (NT). *p<0.05, **p<0.01, ***p<0.001 by 2-way ANOVA test followed by Tukey’s post hoc test. #p<0.001 denotes the statistical comparison between +aWT condition treated with anti-IL-6 antibody and untreated (NT) or IgG-treated +aWT neurons. Analyses were performed on n>74 neurons per experimental group from N>5 biological replicates, except for IgG isotypic treated neurons (n>15), which derived from N=1-3 replicates.

Since the presence of serum in culture might predispose astrocytes towards a reactive phenotype, *IL-6* levels were assessed in astrocytes directly isolated from a mouse model of RTT. To this aim, by taking advantage of MACS technology, astrocytes were acutely isolated from the cortex of P7 heterozygous (HET) animals and the corresponding WT female littermates at P7 (**Figure 3D**), and qRT-PCR indicated a significant increase of *IL-6* mRNA levels in HET astrocytes (**Figure 3E**). Interestingly, Pearson correlation analysis reported that *IL-6* transcriptional levels tend to inversely correlate with the expression of wild-type *Mecp2* (r^2^=0.4341, p=0.075), suggesting that *IL-6* upregulation might be confined to *Mecp2*^-^ astrocytes.

It is known that IL6-dependent signalling induces the activation of the transcription factor Stat3, whose phosphorylation leads to several genomic effects in different cell types, including neurons^26^. As proof of the functional role of astrocyte-derived IL-6, Western blot analysis demonstrated a significant increase in Stat3 phosphorylation in neurons cultured with KO glial cells, that was completely prevented by a neutralizing anti-IL-6 antibody (**Figure 3F,G**). To evaluate whether Stat3 activation was causally related to the altered synapse formation observed in neurons exposed to KO astrocytes, anti-IL-6 antibody was added to co-cultures (**Figure 3G**) and synaptic density evaluated by immunofluorescence. We found that the blockade of IL-6 signalling through the neutralizing antibody rescued the number of pre-synaptic terminals, whereas post-synaptic puncta were only partially recovered. Importantly, the same treatment significantly increased the number of colocalized puncta, therefore restoring functional synapses. The specific effect of the blocking antibody against IL-6 was assessed by testing an isotypic IgG antibody, which did not produce the same beneficial effects. In line with the physiological role of IL-6 in synaptic formation and maintenance^27^, blocking IL-6 in neurons cultured with WT astrocytes reduced the density of post-synaptic puncta (**Figure 3H-K**).

### The secretome of *Mecp2* KO astrocytes exerts synaptotoxic effects independent from IL-6

To understand whether KO astrocytes intrinsically secrete synaptotoxic factors, we we explored the effects of KO ACM on synapses and assessed whether KO astrocytes cultured alone secrete immune-derived signals similar to those released in co-culture with neurons. Unlike the transwell-based co-culture system used so far, WT neurons matured alone and were exposed from DIV13 to DIV14 to ACM, whose composition depends on astrocyte cell-autonomous properties (**Figure 4A**). Analysis of puncta density indicated that neurons treated with WT ACM show a healthy phenotype in terms of pre- and post-synaptic puncta number, with a trend towards an increase compared with neurons treated with the control medium, in line with the well-known pro-synaptogenic properties of ACM (**Figure 4B-E**). Similar to the co-culture condition (**Figure 1**), exposure of WT neurons to KO ACM caused a detrimental effect on both pre- and post-synaptic compartments and their colocalization. In addition, to explore the nature of the causative molecules, KO ACM was heated to denature proteins; the lack of any detrimental effect in neurons treated with heat-inactivated KO ACM (KO ACM*) demonstrated that secreted neurotoxic factors are thermolabile (**Figure 4B-E**).

**Figure 4.**
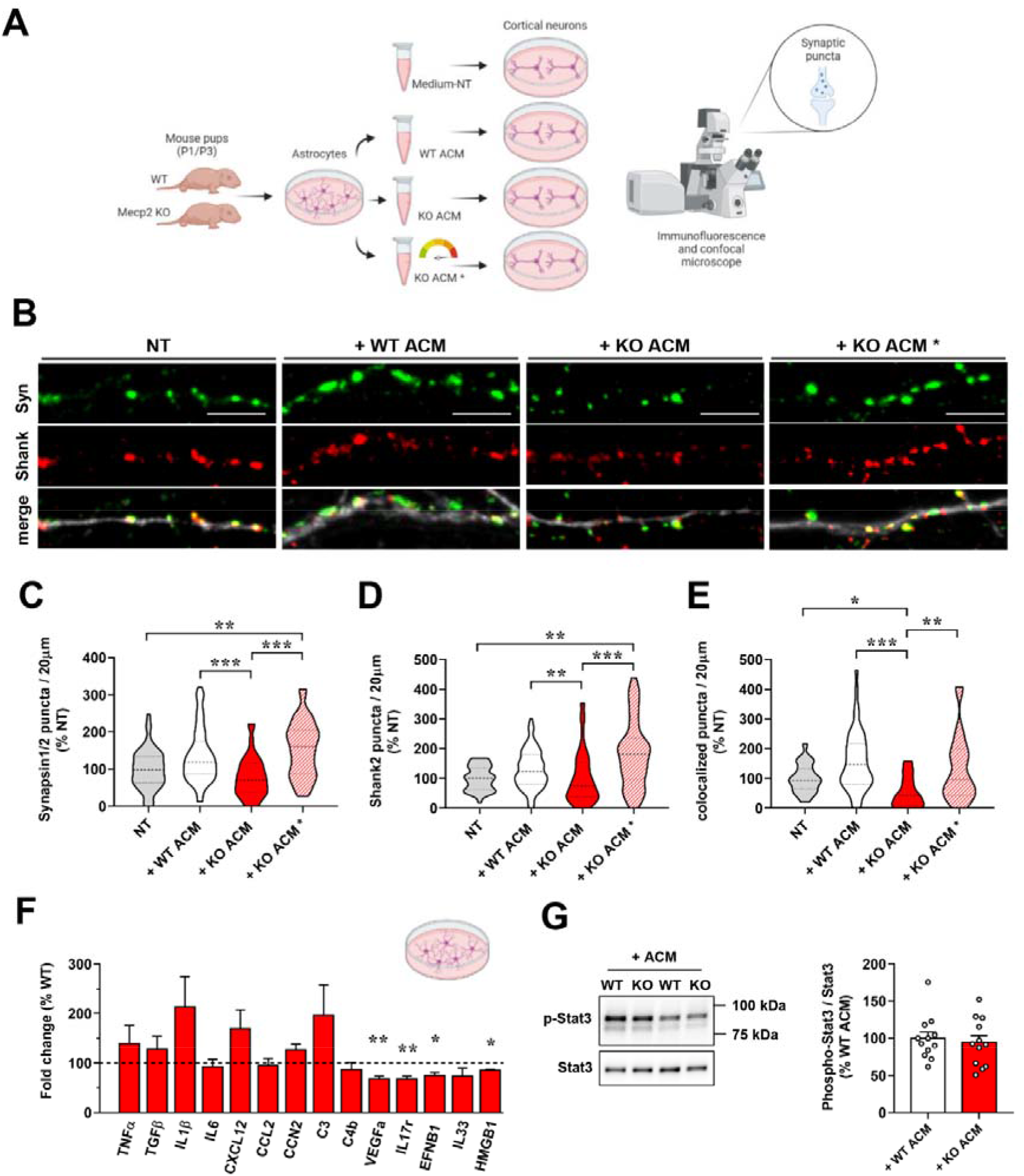
*Mecp2* KO astrocytes cultured alone secrete thermolabile synaptotoxic factors, with no upregulation of IL-6. **(A)** Experimental design overview, created withBioRender.com. **(B)** Representative images of primary branches from WT neurons (DIV14) immunostained for Synapsin1/2 (green), Shank2 (red) and their merge with MAP2 (white). KO ACM* indicates heat-inactivated ACM. Scale bar = 5 μm. **(C-E)** Violin plots indicate the median (dashed line) and 25th and 75th percentiles (dotted lines) of Synapsin1/2 (C), Shank2 (D) and colocalized puncta number (E). Values for puncta number are expressed as percentages of neurons treated with empty medium (NT). *p<0.05, **p<0.01, ***p<0.001 by Kruskal-Wallis test followed by Dunn’s post hoc test. Analyses were performed on n>24 neurons per experimental group from N>4 biological replicates. All data derived from at least 2 independent experiments. **(F)**. The graph depicts the mRNA levels of selected astrocyte genes in *Mecp2* KO cortical astrocytes cultured alone, expressed as percentages of WT astrocytes (n=10). Data are represented as mean±SEM. *p<0.05, **p<0.01 by Student’s t-test or Mann–Whitney test in accordance with data distribution. **(G)** Western blot analysis of phospho-Stat3 over Stat3 protein levels in neurons exposed to WT or KO ACM. Data, represented as mean ±SEM, are expressed as percentages of neurons treated with WT ACM (n=12); representative bands for phosphorylated and total Stat3 are reported.

To understand whether the ACM-induced effects rely on molecular mechanisms similar to those of the co-cultures, we analysed the expression of the same set of pro-inflammatory genes in KO astrocytes cultivated alone. qRT-PCR revealed that mRNA levels of *VEGF*, *IL17R*, *EFNB1*, *IL33*, and *HMGB1* were downregulated; interestingly, the expression of *IL-6*, as well as that of *IL-1β* and *CXCL12,* was not affected (**Figure 4F**). Accordingly, the lack of Stat3 activation in neurons treated with KO ACM excluded the involvement of IL-6 in the ACM-induced synaptic defects (**Figure 4G**).

Taken together, these data demonstrate that *Mecp2* KO astrocytes release synaptotoxic and thermolabile factors by cell-autonomous mechanisms that however differ from IL-6.

## Discussion

In this study we investigated the effect of astrocyte-neuron communication on the capacity of *Mecp2* KO astrocytes to regulate synapse formation via paracrine signals. In agreement with literature, our results confirmed the damaging action exerted by *Mecp2* KO astrocytes on neuronal health, reinforcing the importance of *Mecp2* expression in astrocytes to sustain neuronal maturation by non-cell autonomous mechanisms^16,18,28^. However, by analyzing the density of synaptic puncta, which is profoundly compromised in RTT^13–15^, we demonstrated that Mecp2 KO astrocytes negatively affect synaptogenesis. Further, we disclosed that molecules secreted by KO astrocytes activate in WT neurons an abnormal inflammatory response, triggered by IL-6. Indeed, IL-6 blockade restored synaptic alterations therefore proving the detrimental action of this cytokine, whose aberrant expression in astrocytes requires a crosstalk with neurons.

Several lines of research highlighted the heterogeneity of astrocyte populations and how astrocytes from different brain areas exert distinct effects on neuronal functions^29^. Our results indicated cortical KO astrocytes as crucial cells responsible for the altered synapse density in RTT and this evidence is in line with our recent data demonstrating that cortical glial cells show the most severe cytoskeletal and transcriptional alterations in *Mecp2* KO brain^30^. Notably, although the heterotypic co-cultures in which astrocytes derived from different regions are cultured with cortical neurons allowed to compare the synaptogenic potential of region-specific astrocytes^24^, we cannot exclude that this setting could have masked other alterations proper of the respective homotypic culture.

Diversely from previous studies examining the molecular phenotypes of KO astrocytes alone^19,20,31^, to identify the molecules involved in the occurrence of synaptic defects, we used an alternative strategy aimed at assessing deregulated pathways in neurons cultured with astrocytes to deduce upstream regulators. The validity of this approach was supported by the significant downregulation of pathways associated with synaptic maturation and functionality, in line with our immunofluorescence results. Further, the observation that synapse-related pathways were downregulated also comparing neurons cultured with KO astrocytes to neurons alone suggested that KO astrocytes release synaptotoxic molecules, although a defective secretion of synaptogenic cues cannot be excluded. Indeed, we believe that a synergic cooperation between reduced secretion of synaptogenic factors and increased release of toxic molecules might contribute to the observed synaptic defects.

The novelty of our study also relies on the evidence that the neuronal-astrocyte crosstalk impinges on the *Mecp2* KO astrocyte secretome. Indeed, our molecular data indicated a significant upregulation of IL-6 in KO astrocytes only when maintained in communication with neurons. In agreement, transcriptomic and proteomic studies performed in KO astrocytes and their medium did not report an increase of IL-6 so far^19,20,31^, underlying the importance of astrocyte-neuron communication for dictating the molecular and functional properties of astrocytes. The upregulation of IL-6 in astrocytes isolated from the cortex of P7 heterozygous animals strengthened its involvement in RTT and excluded the possibility of methodological artifacts of culturing cells in a medium supplemented with serum. In addition, although in the present work we have not discriminated *Mecp2^+^* from *Mecp2^-^*astrocytes, the trend of an inverse correlation between *IL-6* and *Mecp2* expressions suggests an increased production of the cytokine by astrocytes expressing the mutant allele.

IL-6 is a cytokine involved in the regulation of neuronal and synaptic functions^27^, and, as occurs for this family of proteins, it exhibits pleiotropic actions depending on its concentration, displaying either neurotrophic properties or detrimental activity^32^. Indeed, contrasting data are available regarding the effects of IL-6 on excitatory synapse formation^27^, which might depend on the source of IL-6 and its concentration. Our results support the importance of a fine regulation of its levels since decreasing IL-6 levels in WT co-cultures proved to be detrimental to post-synaptic terminals. Conversely, the same blocking strategy in neurons cultured with KO astrocytes, therefore exposed to excessive IL-6 levels, was effective in reducing IL-6 mediated pathway and improving synaptic phenotypes.

The relevance of our findings is emphasized by several evidences. First, numerous clinical and experimental data point to the presence of a subclinical inflammatory status in RTT, characterized by cytokine dysregulation and aberrant NF-kB pathway^33–37^. Of note, IL-6 increased expression has already been described in the brain, saliva and plasma of patients suffering from RTT^34,38^, as well as in other neuropsychiatric disorders^39^. Further, IL-6 overexpression in the mouse brain correlates with neurological abnormalities^40,41^ and its inhibition improves social behaviors^42^. Accordingly, IL-6 and its pathway have been already proposed for the development of novel therapies in mental diseases, which however should aim at attenuating IL-6 activity without completely abolishing it, considering the above-mentioned physiological roles^42,43^.

In conclusion, our study identified IL-6 as a non-cell autonomous synaptotoxic mechanism triggered by *Mecp2* KO astrocytes, providing an interesting therapeutic target for RTT and other *MECP2*-related disorders.

## Acknowledgments

Part of this work was carried out at NOLIMITS, an advanced imaging facility established by the Università degli Studi di Milano. RNA-Seq analysis was performed by GENARTIS srl, with the support of Dr. Marzia Rossato and Dr. Giulia Lopatriello. Flow cytometry analyses were performed by the Flow Citometry Unit (FRACTAL) at San Raffaele Scientific Institute, Milan. We are grateful to all members of AF and NL laboratories for helpful discussions. This work was mainly supported by the Italian parents’ association “Pro RETT Ricerca” to AF and NL.

## Author contributions

EA, EF, MB, CC and AF conducted the experiments and data analysis. EA and AF designed the experiments, prepared the figures and wrote the manuscript. DP provided his expertise in IL-6 mediated signalling, supporting analysis of IL-6 signalling and experiments with anti-IL-6 antibody. EB provided technical support for MACS Technology. CB supported the bioinformatics analyses. CDP contributed to molecularly validate data obtained from transcriptomic analyses. CDQ performed RNA-Seq and bioinformatics analyses. NL revised the manuscript and assisted in planning, interpreting and discussing results.

## Data availability

RNA-seq data in this study have been deposited in the ArrayExpress repository (E-MTAB-12393). They are publicly available as of the date of publication.

## Excel tables

**Table S1.** All significant GO terms from enrichment analysis on downregulated DEGs in +aKO *vs* +aWT comparison (p.adj<0.1).

**Table S2.** All significant GO terms from enrichment analysis on downregulated DEGs in +aKO *vs* CTRL comparison (p.adj<0.1).

**Table S3.** All dysregulated gene sets from GSEA analysis on downregulated DEGs in +aKO *vs* +aWT comparison (FDR q<0.05).

**Table S4.** All significant GO terms from enrichment analysis on upregulated DEGs in +aWT *vs* CTRL comparison (p.adj<0.1).

**Table S5.** All significant GO terms from enrichment analysis on upregulated DEGs in +aKO *vs* CTRL comparison (p.adj<0.1).

**Table S6.** List of primers for genotyping

**Table S7.** List of primers for qRT-PCR

