## Supplementary data for "*Mecp2* knock-out astrocytes affect synaptogenesis by IL-6 dependent mechanisms"

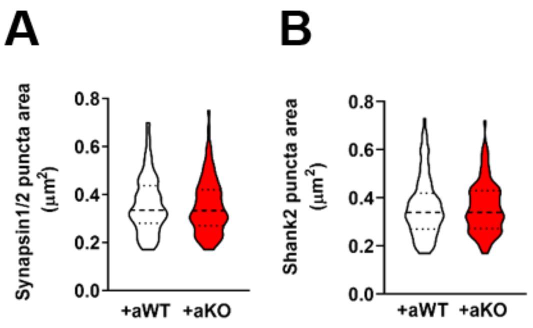


**Suppl. Figure 1. Soluble factors secreted by *Mecp2* KO astrocytes do not affect puncta size**

Violin plot indicates the median (dashed line) and 25th and 75th percentiles (dotted lines) of Synapsin1/2 (A) and Shank2 (B) puncta area of neurons co-cultured with WT or KO cortical astrocytes seeded on transwell inserts. Data are represented as mean±SEM. Analyses were performed on n>107 neurons from N>15 biological replicates. Samples derived from 4 independent experiments.

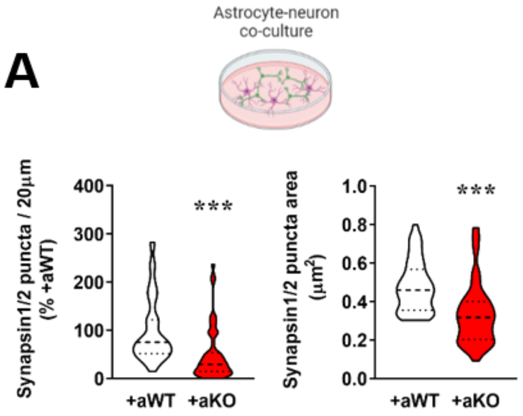


**Suppl. Figure 2. *Mecp2* KO astrocytes induce a significant reduction of pre-synaptic puncta number and size when cultured in contact with WT neurons**

Violin plots indicate the median (dashed line) and 25th and 75th percentiles (dotted lines) of Synapsin1/2 puncta density (B) and puncta area (C) of neurons cultured in contact with WT or KO cortical astrocytes. Data are indicated as mean±SEM. ***p<0.001 by Mann-Whitney test. Analyses were performed on n>49 neurons per experimental group from N=6 biological replicates. Samples derived from 2 independent experiments.

**A**

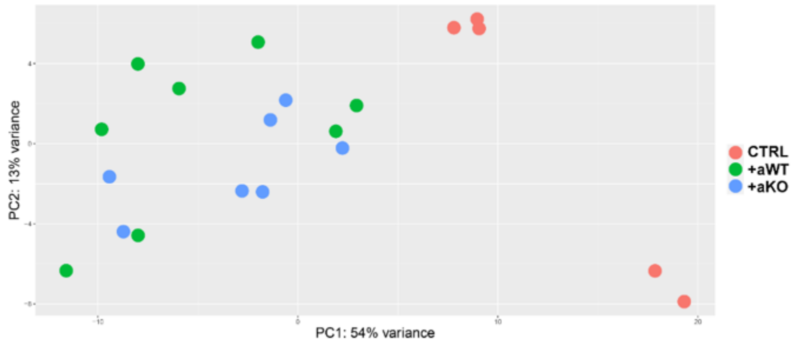


**B**

| Comparison | p.adj<0.1 | p.adj<0.1 LFC>\|1\| | p.adj<0.1 LFC>1 | p.adj<0.1 LFC<-1 | p.adj<0.05 | p.adj<0.05 LFC>\|1\| | p.adj<0.05 LFC>1 | p.adj<0.05 LFC<-1 |
|---|---|---|---|---|---|---|---|---|
| +aWT vs CTRL | 3239 | 545 | 437 | 108 | 2493 | 545 | 437 | 108 |
| +aKO vs CTRL | 3808 | 717 | 573 | 144 | 2890 | 716 | 572 | 144 |
| +aKO vs +aWT | 401 | 25 | 24 | 1 | 216 | 19 | 18 | 1 |

**C**

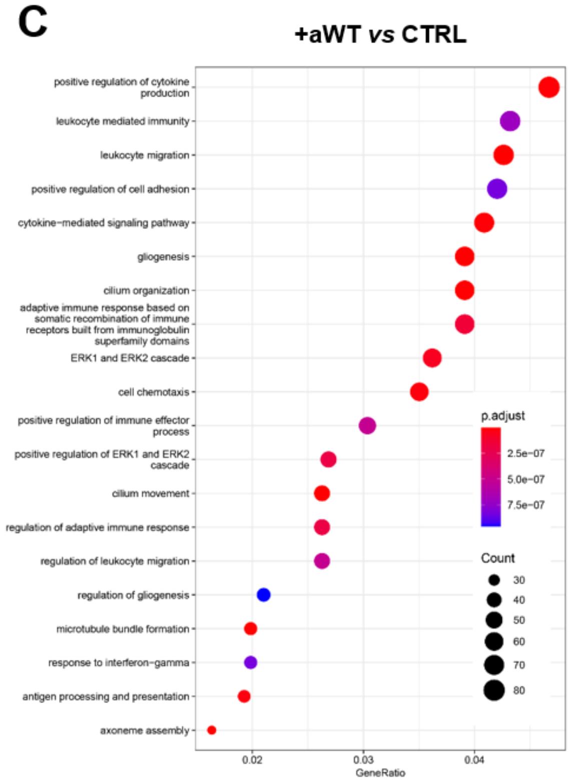


**D**

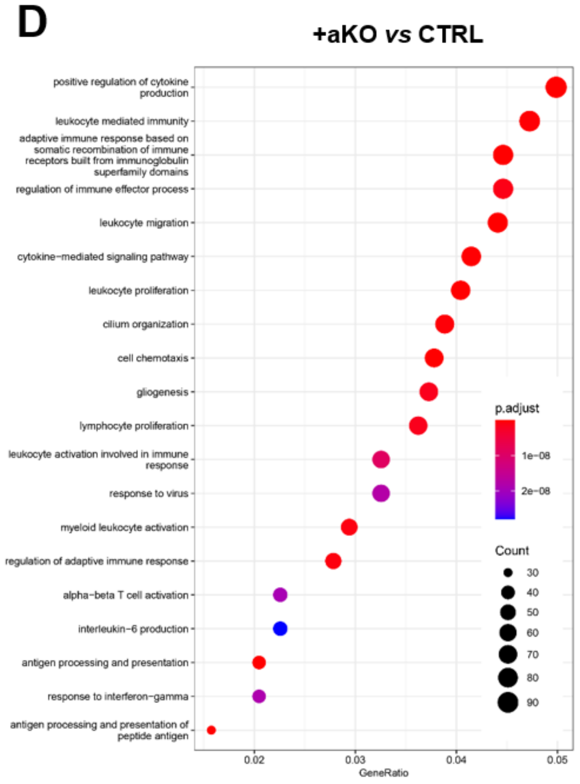


**Suppl. Figure 3**. ***Mecp2* KO astrocytes alter gene expression in WT co-cultured neurons**

**(A)** Principal Component Analysis (PCA) of RNA Seq data. Gene expression was investigated in neurons cultured alone (CTRL), in neurons cultured with WT (+aWT) or *Mecp2* KO (+aKO) astrocytes. PCA was performed using normalized RNA-Seq data. A clear difference was observed between CTRL and +aWT/+aKO samples. **(B)** The table reports the number of differentially expressed genes (DEGs) in the different comparisons, filtering on the basis of FDR adjusted p-value (p.adj) and log2 fold change (LFC). **(C,D)** Enrichment analysis data of +aWT *vs* CTRL (C) and +aKO *vs* CTRL comparisons (D) with upregulated DEGs at p.adj<0.1 (see Table S4 and S5, respectively), showing the top 20 significant GO terms (biological process). Size and colour of each dot represent -$\log_2$ of FDR and number of genes associated with each term, respectively, according to the scale indicated in the figure.

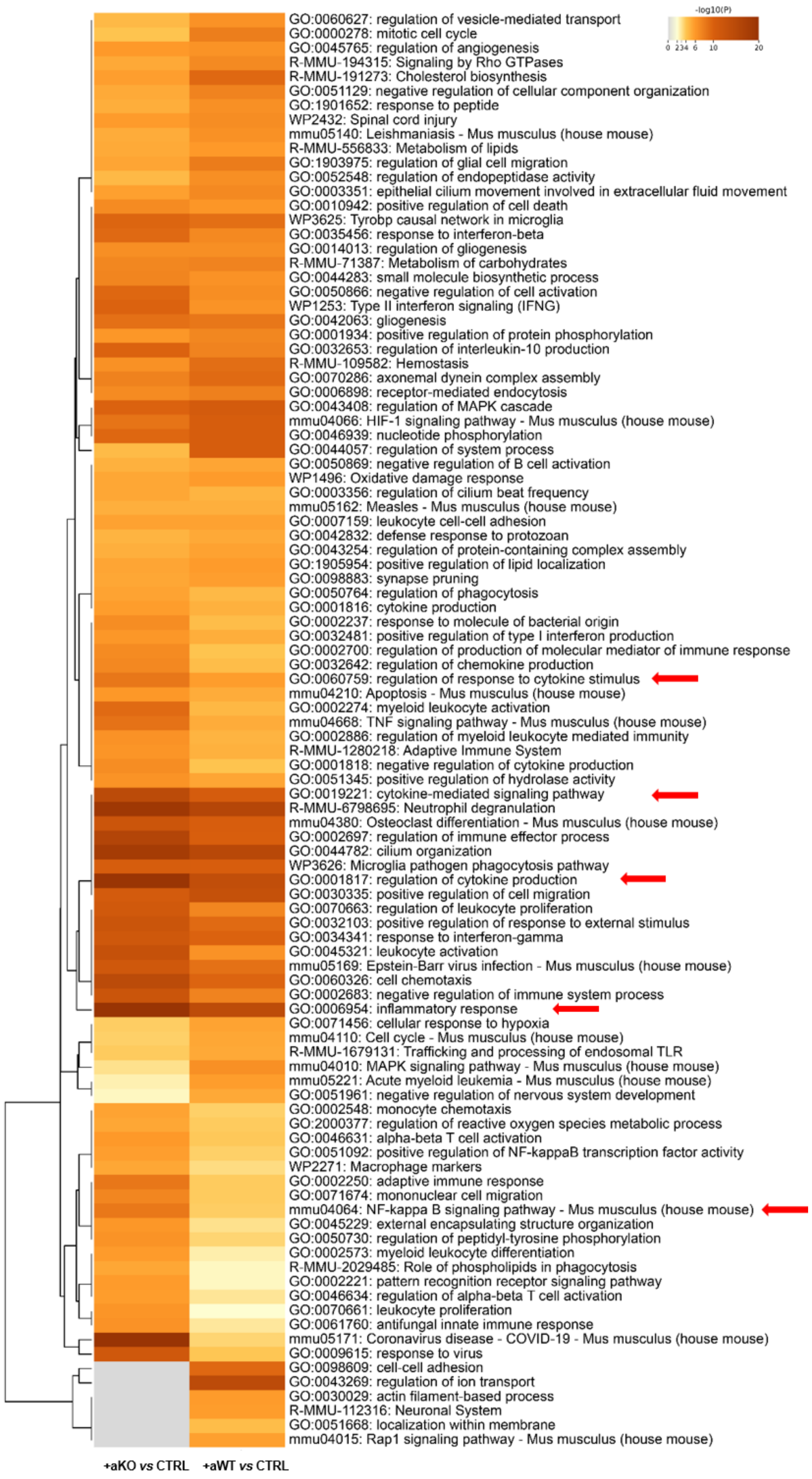
-log10(P)
0 2 3 4 6 10 20
GO:0060627: regulation of vesicle-mediated transport
GO:0000278: mitotic cell cycle
GO:0045765: regulation of angiogenesis
R-MMU-194315: Signaling by Rho GTPases
R-MMU-191273: Cholesterol biosynthesis
GO:0051129: negative regulation of cellular component organization
GO:1901652: response to peptide
WP2432: Spinal cord injury
mmu05140: Leishmaniasis - Mus musculus (house mouse)
R-MMU-556833: Metabolism of lipids
GO:1903975: regulation of glial cell migration
GO:0052548: regulation of endopeptidase activity
GO:0003351: epithelial cilium movement involved in extracellular fluid movement
GO:0010942: positive regulation of cell death
WP3625: Tyrobp causal network in microglia
GO:0035456: response to interferon-beta
GO:0014013: regulation of gliogenesis
R-MMU-71387: Metabolism of carbohydrates
GO:0044283: small molecule biosynthetic process
GO:0050866: negative regulation of cell activation
WP1253: Type II interferon signaling (IFNG)
GO:0042063: gliogenesis
GO:0001934: positive regulation of protein phosphorylation
GO:0032653: regulation of interleukin-10 production
R-MMU-109582: Hemostasis
GO:0070286: axonemal dynein complex assembly
GO:0006898: receptor-mediated endocytosis
GO:0043408: regulation of MAPK cascade
mmu04066: HIF-1 signaling pathway - Mus musculus (house mouse)
GO:0046939: nucleotide phosphorylation
GO:0044057: regulation of system process
GO:0050869: negative regulation of B cell activation
WP1496: Oxidative damage response
GO:0003356: regulation of cilium beat frequency
mmu05162: Measles - Mus musculus (house mouse)
GO:0007159: leukocyte cell-cell adhesion
GO:0042832: defense response to protozoan
GO:0043254: regulation of protein-containing complex assembly
GO:1905954: positive regulation of lipid localization
GO:0098883: synapse pruning
GO:0050764: regulation of phagocytosis
GO:0001816: cytokine production
GO:0002237: response to molecule of bacterial origin
GO:0032481: positive regulation of type I interferon production
GO:0002700: regulation of production of molecular mediator of immune response
GO:0032642: regulation of chemokine production
GO:0060759: regulation of response to cytokine stimulus
mmu04210: Apoptosis - Mus musculus (house mouse)
GO:0002274: myeloid leukocyte activation
mmu04668: TNF signaling pathway - Mus musculus (house mouse)
GO:0002886: regulation of myeloid leukocyte mediated immunity
R-MMU-1280218: Adaptive Immune System
GO:0001818: negative regulation of cytokine production
GO:0051345: positive regulation of hydrolase activity
GO:0019221: cytokine-mediated signaling pathway
R-MMU-6798695: Neutrophil degranulation
mmu04380: Osteoclast differentiation - Mus musculus (house mouse)
GO:0002697: regulation of immune effector process
GO:0044782: cilium organization
WP3626: Microglia pathogen phagocytosis pathway
GO:0001817: regulation of cytokine production
GO:0030335: positive regulation of cell migration
GO:0070663: regulation of leukocyte proliferation
GO:0032103: positive regulation of response to external stimulus
GO:0034341: response to interferon-gamma
GO:0045321: leukocyte activation
mmu05169: Epstein-Barr virus infection - Mus musculus (house mouse)
GO:0060326: cell chemotaxis
GO:0002683: negative regulation of immune system process
GO:0006954: inflammatory response
GO:0071456: cellular response to hypoxia
mmu04110: Cell cycle - Mus musculus (house mouse)
R-MMU-1679131: Trafficking and processing of endosomal TLR
mmu04010: MAPK signaling pathway - Mus musculus (house mouse)
mmu05221: Acute myeloid leukemia - Mus musculus (house mouse)
GO:0051961: negative regulation of nervous system development
GO:0002548: monocyte chemotaxis
GO:2000377: regulation of reactive oxygen species metabolic process
GO:0046631: alpha-beta T cell activation
GO:0051092: positive regulation of NF-kappaB transcription factor activity
WP2271: Macrophage markers
GO:0002250: adaptive immune response
GO:0071674: mononuclear cell migration
mmu04064: NF-kappa B signaling pathway - Mus musculus (house mouse)
GO:0045229: external encapsulating structure organization
GO:0050730: regulation of peptidyl-tyrosine phosphorylation
GO:0002573: myeloid leukocyte differentiation
R-MMU-2029485: Role of phospholipids in phagocytosis
GO:0002221: pattern recognition receptor signaling pathway
GO:0046634: regulation of alpha-beta T cell activation
GO:0070661: leukocyte proliferation
GO:0061760: antifungal innate immune response
mmu05171: Coronavirus disease - COVID-19 - Mus musculus (house mouse)
GO:0009615: response to virus
GO:0098609: cell-cell adhesion
GO:0043269: regulation of ion transport
GO:0030029: actin filament-based process
R-MMU-112316: Neuronal System
GO:0051668: localization within membrane
mmu04015: Rap1 signaling pathway - Mus musculus (house mouse)
+aKO vs CTRL
+aWT vs CTRL

**Suppl. Figure 4. Metascape analysis on upregulated DEGs indicates that pathways related to inflammation are more represented in the +aKO *vs* CTRL with respect to the +aWT *vs* CTRL comparison.**

Heatmap of top 100 enriched terms across the two gene lists, colored by p values.
